# Aggregation of biological particles under radial directional guidance

**DOI:** 10.1101/096602

**Authors:** Ion Bica, Thomas Hillen, Kevin J. Painter

## Abstract

Many biological environments display an almost radially-symmetric structure, allowing proteins, cells or animals to move in an oriented fashion. Motivated by specific examples of cell movement in tissues, pigment protein movement in pigment cells and animal movement near watering holes, we consider a class of radially-symmetric anisotropic diffusion problems, which we call the *star problem*. The corresponding diffusion tensor *D*(*x*) is radially symmetric with isotropic diffusion at the origin. We show that the anisotropic geometry of the environment can lead to strong aggregations and blow-up at the origin. We classify the nature of aggregation and blow-up solutions and provide corresponding numerical simulations. A surprising element of this strong aggregation mechanism is that it is entirely based on geometry and does not derive from chemotaxis, adhesion or other well known aggregating mechanisms. We use these aggregate solutions to discuss the process of pigmentation changes in animals, cancer invasion in an oriented fibrous habitat (such as collagen fibres), and sheep distributions around watering holes.

**JTB classification:** 21.050, 21.160, 52.250, 71.060

## 1. Introduction

Movement of biological particles, whether molecules, cells or organisms, is heavily dictated by their environment. We explore the impact of oriented environments, where a particle’s motion is influenced by the anisotropic nature of its surroundings. Examples cover a broad range of biological scales: within single cells, the structured cytoskeleton offers a transport system for efficient shuttling of molecules and organelles [1]; in tissues, cell migration and consequently tumour invasion can be facilitated by movement along collagen fibres, neuronal axons and capillaries [9]; at a landscape level, animals often follow (or avoid) paths, roads and other linear structures [16, 4, 21]. Oriented environments may also have less manifestly physical forms: chemicals, the geomagnetic field, sound, visual cues and many other factors can present orientating information.

In [15] we termed the *fully anisotropic diffusion equation* as the linear parabolic equation

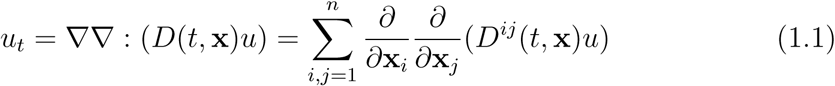

on a bounded or unbounded domain in ℝ^*n*^, equipped with appropriate boundary conditions. The tensor *D*(*t*, **x**) = (*D^ij^*(*t*, **x**))_*i,j*_ describes anisotropic diffusion, in which diffusive spread is distinct along different axial directions. This model arises as a description for particle movement in terms of their macroscopic density *u*(*t*, **x**): models similar to (1.1) have been used to explain cell migration along collagen fibres (see [11]) and the invasion of glioma (brain tumour) cells along neural fibre tracts (see [28, 6]); for animal populations, they have been used to describe wolf movement along linear features in boreal habitats [14, 20, 21], sea turtle navigation [29] and butterfly movement [27]. The wolf movement problem led to the specific exploration into the dynamics of (1.1) under a mathematically convenient straight-line structure, such as a road: the population is shown to accumulate onto the line and, under certain limiting scenarios, solutions blow up in infinite time [15].

A second logical abstraction is to assume radially-aligned orienteering information. With respect to earlier examples, cytoskeletal microtubules are arranged into spokes radiating from the cell nucleus (or other organising centre) [1]; radially-aligned collagen fibres can be found in both healthy and pathological scenarios, such as emanating from the nipple of mammary tissue or oriented orthogonally to the tumour boundary in malignant breast tumours [30]; at a landscape scale, animal trails radiate from waterholes in arid environments [18]. The *star problem* investigated here emerges as an idealised description of movement along (or perpendicular to) radial lines that converge on some origin, under assumed radial symmetry. Given a planar polar coordinate system (*r*, *ϕ*), where for each point the direction away from the origin will be given by radial unit vector (cos *ϕ*, sin *ϕ*), and assuming that movement is aligned radially, the fully anisotropic diffusion problem (1.1) is shown to have diffusion tensor

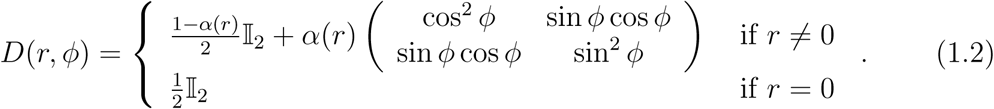

For now we simply state |*α*(*r*) | ≤ 1 to be a given function that describes the precision to which movement along the radial direction (positive *α*) or perpendicular to it (negative *α*) is maintained. Radially symmetric solutions to (1.1) under (1.2) are then found to be determined by the star problem,

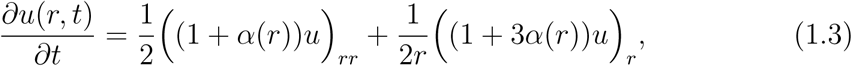

on the interval (0, *R*] (where *R* is potentially infinity), under suitable boundary and initial conditions.

**Figure 1:**
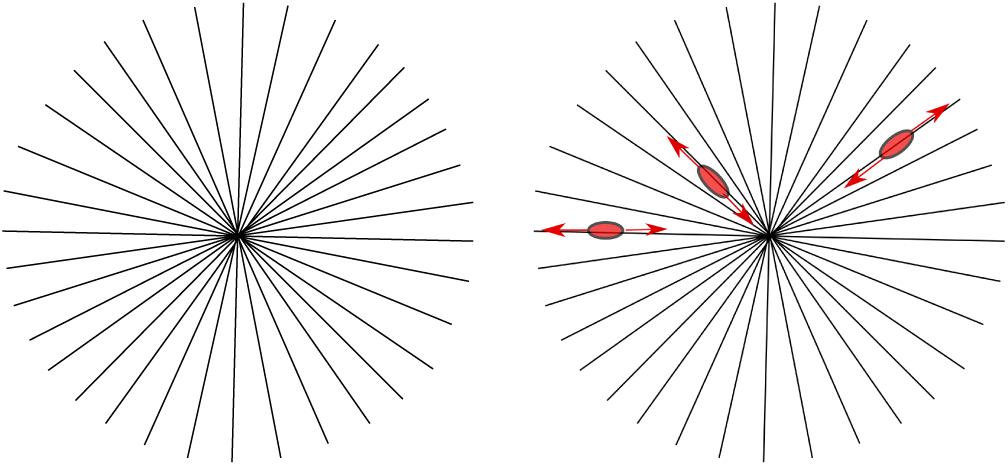
Left: Schematic of the anisotropy of the star problem. Right: Sketch of moving particles in the star domain.

We explore whether the equation (1.3) is capable of creating aggregative behaviour, whether blow-up is possible and whether this blow-up occurs in finite or infinite time. For 0 ≤ *α* ≤ 1 we show that solutions to equation (1.3) contain a leading order term of the form

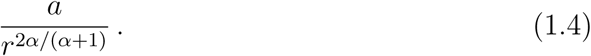

In particular, when *α* = 0 we obtain constant solutions and for *α* = 1 we observe a 1/*r* singularity at 0. The latter implies that solutions to the star problem have the potential to instantly blow up, representing a “strong aggregation” at the origin. This arises purely from the underlying structure of the environment and is hence distinct from the typical aggregations associated with a process such as chemotaxis. For orientation along radial circles (perpendicular to radial directions, −1 ≤ *α* < 0) aggregation does not occur and solutions to (1.3) remain bounded for all time.

### 1.1. Outline

In the following Section 2 we systematically motivate equation (1.3) as an idealised description for movement of biological particles in radially symmetric oriented environments. Starting with a transport equation for the aligned movement of particles, we state the macroscopic continuous drift-anisotropic diffusion equation obtained under scaling. Equation (1.3) subsequently emerges following the transformation to polar coordinates and assuming radial symmetry. Specific examples are provided within the context of organelle transport along microtubules, cell movement along collagen fibres and animal movement along the trails that surround water holes. For the special case of constant *α*, the resulting singular Sturm-Liouville problem is solved in Subsection 3.3. The leading order term in this solution is derived and we use Section 4 to compare numerical solutions of (1.2) to the asymptotic formula, as well as demonstrate the utility of the model in applications. We conclude with a discussion of the results in the context of our motivating examples.

## 2. Derivation and motivations

Velocity-jump random walk models [24] describe movement as a piecewise-continuous path of smooth runs punctuated by turns into new velocities. As such they provide a plausible approximation of actual movement paths and can be parametrised against standard datasets. The transport equation is the corresponding continuous description for this process and, in a series of papers ([13, 25, 11, 14]), it is shown how first choosing a distribution where turns are biased into specific axial directions and then taking a course-grain limit leads to the anisotropic diffusion formulation (1.1), with appropriate diffusion tensor *D*.

### 2.1. Transport equations to anisotropic diffusion

The transport equation postulates the time evolution of the particle population distribution, *p*(*t*, **x**, **v**), parametrised by time *t* ∈ [0, ∞), position **x** ∈ Ω ⊂ ℝ^*n*^ and velocity **v** ∈ *V* ⊂ ℝ^*n*^. Here, turning is chosen to be (effectively) instantaneous and the new velocity is assumed to not depend on the previous velocity, yielding

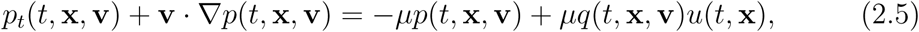

where *u*(*t*, **x**) = ʃ_V_ *p*(*t*, **x**, **v**)*d***v** is the macroscopic (or observable) particle density. *q*(*t*, **x**, **v**) describes the turning distribution, i.e. the probability that a particle turns into velocity **v** at time *t* and position **x**. The parameter *μ* measures the turning rate and is taken here to be constant. We also simplify by assuming that the particles move with a fixed mean speed *s* and, consequently, trivial rescaling allows us to set *s* = *μ* = 1 (and subsequently drop these notations). Hence *V* ≡ 𝕊^n–1^ (the unit sphere) and *q*(*t*, **x**, **v**) becomes a directional distribution on the unit sphere,

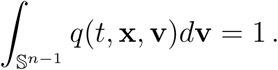

In fact, an equation similar in form to (2.5) was used in [26] to model cell migration along collagen fibres. There, radial fibre arrays were shown to generate focussed aggregations at the origin, suggesting that oriented environments could act to spatially organise populations. Those simulations partially motivate the current work, where a more detailed investigation is conducted through exploring the macroscopic version. This macroscopic model can be obtained through moment closure techniques

(see [14]), creating a drift-anisotropic diffusion equation of the form

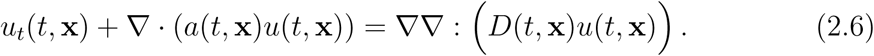

The advective velocity **a**(*t*, **x**) and the anisotropic diffusion tensor *D*(*t*, **x**) derive from statistical properties of the turning distribution:

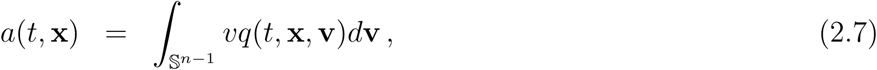

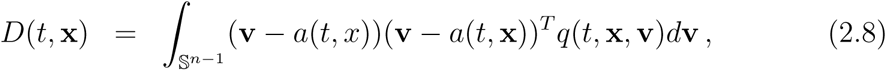

which are respectively the expectation and variance-covariance matrix of *q*. It is noted that the diffusion tensor *D* is symmetric.

Symmetric (or bidirectional) *q*, i.e. *q*(*t*, **x**, **v**) = *q*(*t*, **x**, −**v**) result in the advective velocity *a* vanishing and we obtain precisely the fully anisotropic-diffusion model (1.1), along with anisotropic diffusion tensor

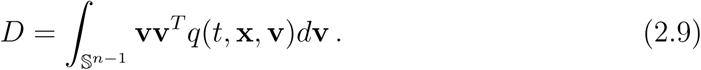

It is noted that an identical model can be obtained under a parabolic scaling (see [14]).

#### 2.1.1. Turning distribution

For the remainder of this paper we restrict to two dimensions, i.e. *n* = 2. In the context of earlier motivating examples, animal movement across landscapes is naturally two-dimensional in scope. Cells are, of course, three-dimensional objects, but studied under *in vitro* conditions can adopt a flattened morphology with cellular structure predominantly extended across the plane (e.g. [7]). Similarly, while most tumours are complicated three-dimensional structures, *in vitro* invasion assays often impose a quasi two-dimensional arrangement (e.g. [31]).

In 2D choosing *q* corresponds to selecting an appropriate directional distribution on the unit circle. Given its status as a circular analogue to the Normal distribution, the von Mises distribution (e.g. [19]) occupies a prominent position for directional datasets and is given by

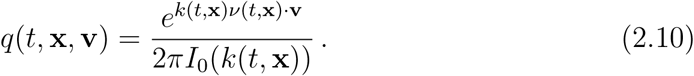

Functions *k*(*t*, **x**) and *ν* (*t*, **x**) define a concentration parameter and dominant direction respectively, while *I_j_*(*k*(*t*, **x**)) denotes the modified Bessel function of order *j*; we refer to Figure 2 for illustrations of (2.10). Notable limits include *k* → 0, generating a uniform distribution, and *k* → ∞, generating a singular distribution that ensures selection of the dominant direction *ν*. Substituting (2.10) into (2.7–2.8) and integrating gives

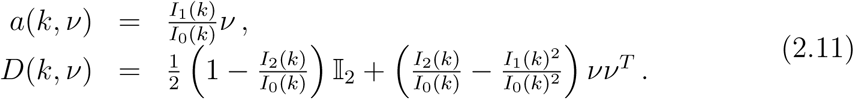

**Figure 2:**
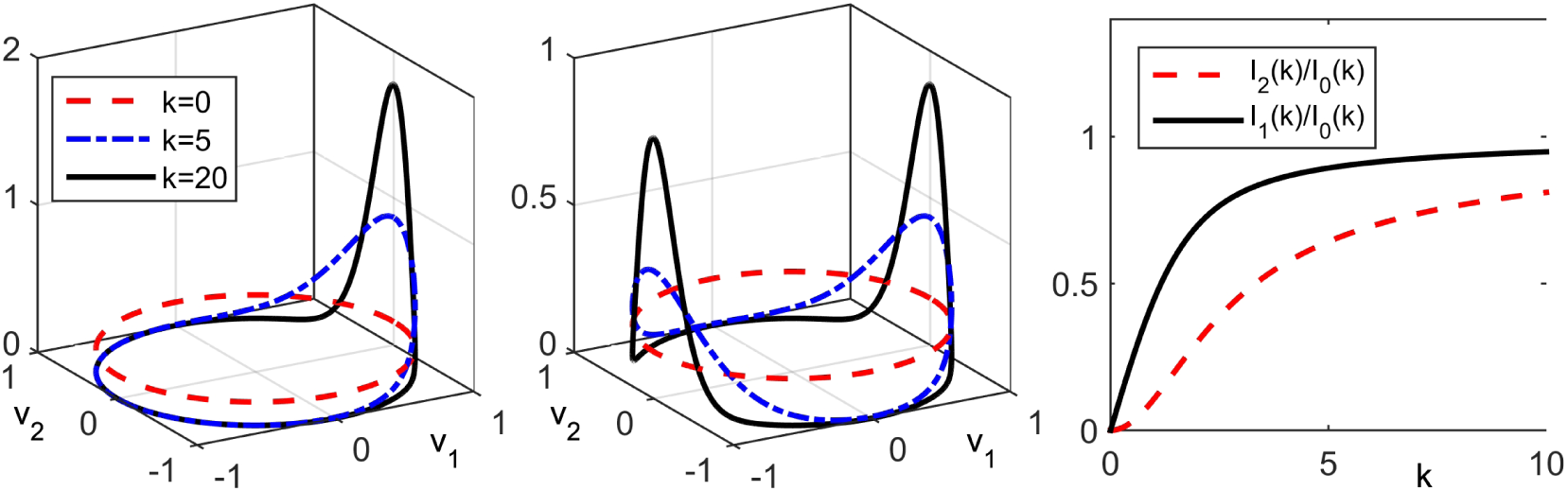
(a) Plot of unidirectional von Mises distribution, *q*(**v**), for *ν* = (1,0) and different values of *k*. (b) Plot of bidirectional von Mises distribution for *ν* = (1,0) and different values of *k*. (c) Plots of *I*_1_(*k*)/*I*_0_(*k*) and *I*_2_(*k*)/*I*_0_(*k*) as functions of *k*.

Note that 𝕀_2_ is the 2 × 2 identity matrix. The functions *I*_1_(*k*)/*I*_0_(*k*) and *I*_2_(*k*)/*I*_0_(*k*) have the following properties:

- *I*_1_(0)/*I*_0_(0) = *I*_2_(0)/*I*_0_(0) = 0;
- they are monotonically increasing for *k* > 0;
- lim_*k*→∞_ *I*_1_(*k*)/*I*_0_(*k*) = lim_*k*→∞_ *I*_2_(*k*)/*I*_0_(*k*) = 1.

Furthermore, we remark that *I*_2_(*k*)/*I*_0_(*k*) < *I*_1_(*k*)^2^/*I*_0_(*k*)^2^ < *I*_1_(*k*)*I*_0_(*k*) < 1; we plot these functions in Figure 2 (c).

The von Mises distribution easily extends to multimodal form through linear combination. Particularly relevant here is the symmetric case of *bidirectional turning*, where turns occur equally into directions *ν* or −*ν*. Such distributions could apply, for example, to animal movement along a path when no particular direction along the path is favoured. In this case,

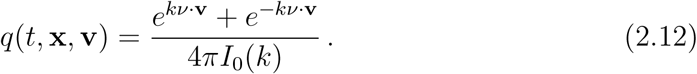

The symmetry in *q* ensures *a* = 0 while

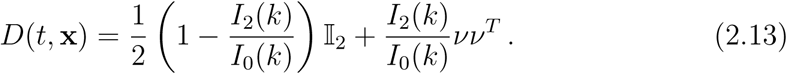

For *k* = 0 we obtain a uniform distribution and recover isotropic diffusion, while for *k* → ∞ we obtain “fully anisotropic” diffusion, where diffusion only takes place along the axis of the dominant direction.

#### 2.1.2. Radial alignment

In a radially symmetric environment we assume the dominant movement direction follows radial lines that converge on some origin. With the bimodal von Mises distribution we choose *ν*(**x**) = ±**x**/ |**x**| for the dominant direction. In this case it is natural to adopt planar polar coordinates **x** = (*r* cos *ϕ*, *r* sin *ϕ*), with radial coordinate *r* ≥ 0 and angular coordinate *ϕ* ∈ [0, 2π]. Therefore, *ν* = ±(cos *ϕ*, sin *ϕ*)^T^. Variation in the concentration parameter is still possible, however we restrict to assuming *k* varies only with distance from the origin (i.e. *k* = *k*(*r*)). Under bidirectional turning we take *D* from equation (2.13) and, substituting *ν* and *k* and setting *α*(*r*) = *I*_2_(*k*(*r*))/*I*_0_(*k*(*r*)) ∈ [0,1)), obtain equation (1.1) along with (1.2) for the macroscopic movement model.

As a further remark, the same procedure applies under radially symmetric cases where the dominant movement direction is orthogonal to radial lines; relevant scenarios include tumour encapsulation by collagen with a predominant fibril orientation parallel to the tumour surface (e.g. [30]). Under this situation the dominant direction of movement is given by *ν*^┴^ = (sin *ϕ*, − cos *ϕ*)^T^. We define a bimodal von-Mises distribution in direction ±*ν*^┴^ and find the diffusion tensor (2.13) to become

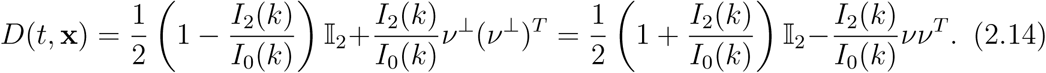

Hence the coefficient *I*_2_(*k*(*r*))/*I*_0_(*k*(*r*)) appears with an opposite sign compared to (2.13).

#### 2.1.3. Transformation into polar coordinates

To transform equation (1.1) into polar coordinates we begin with a general symmetric diffusion tensor

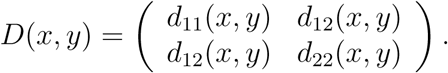

Equation (1.1) in 2D becomes

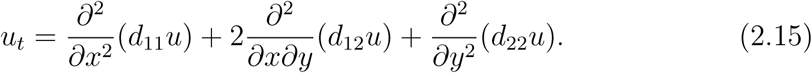

In polar coordinates we have for *r* > 0 that

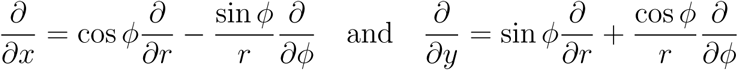

and repeated application of these derivatives gives the second order terms:

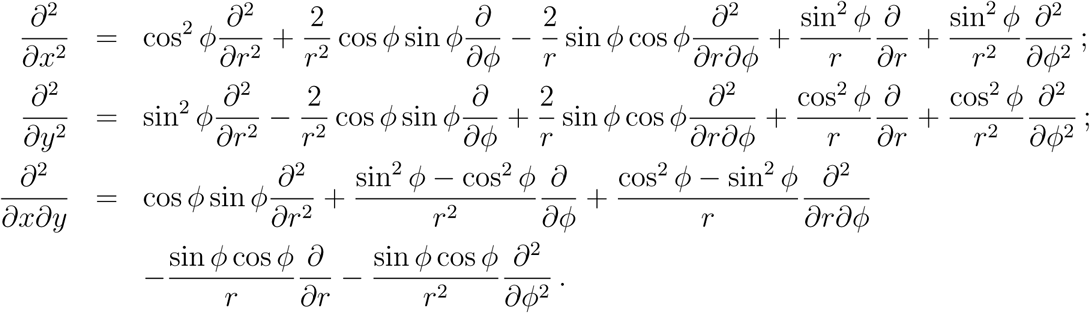

For the symmetric choice *D*(*x*,*y*) = *D*(*ϕ*) we look for radially symmetric solutions *u*(*r*,*t*). Then the terms in the fully anisotropic diffusion model become:

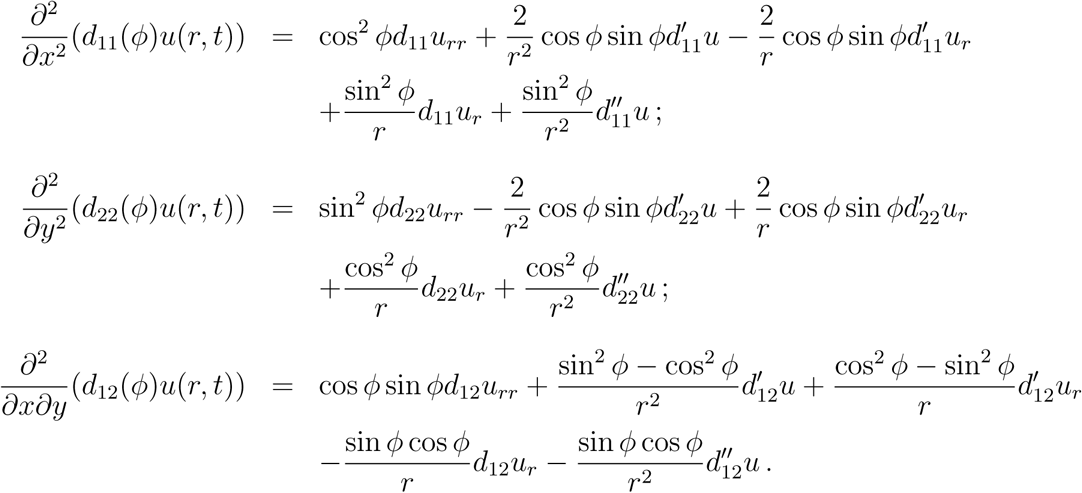

Extending to the choice of *D* in (1.2) we find coefficients

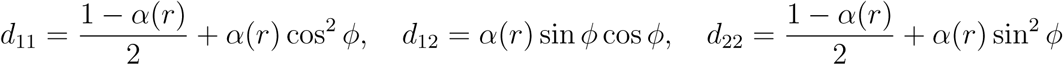

and obtain:

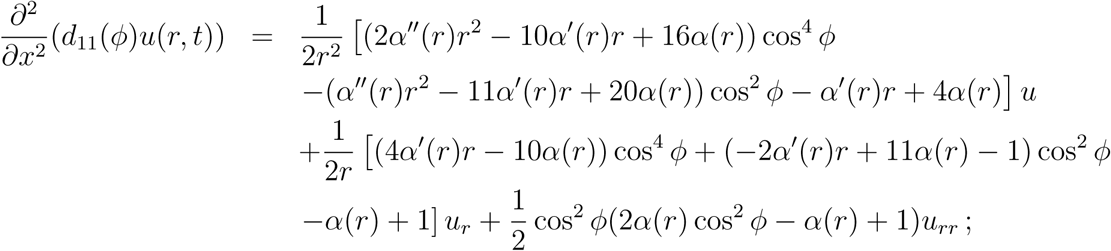

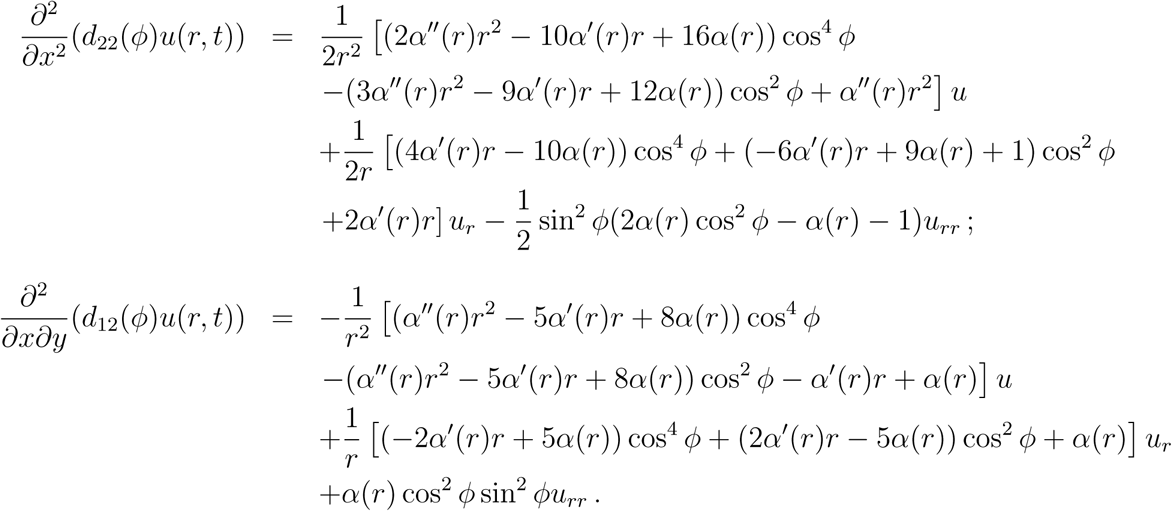

Adding these terms according to (2.15) and simplifying yields the star problem

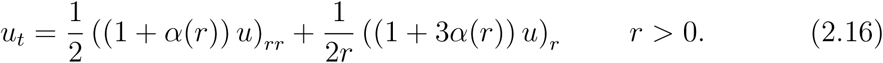

Two special cases are noted as follows:

- For *α*(*r*) = 1 we obtain

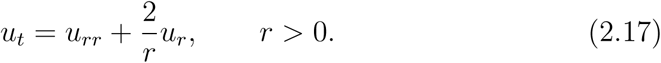
- For *α*(*r*) = 0 we obtain

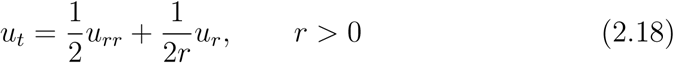

which simply corresponds to 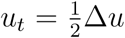 under radially symmetric solutions in planar polar coordinates.

The corresponding boundary conditions of these problems will be discussed later. They vary between the different applications.

### 2.2. Biological examples

In this section we consider the star problem in the context of a special case for three distinct movement processes: melanosome transport during intracellular trafficking, cell movement in extracellular matrix and animal movement around watering holes. These diverse applications are intended to demonstrate the broadness of the star problem across different processes and the modelling itself is kept intentionally simple to transparently connect the application to the more abstract problem.

#### 2.2.1. Melanosome transport

Intracellular shuttling of molecules and organelles is key for many cell functions. The movement of melanosomes (organelles that synthesise and deposit the pigment melanin) in pigment cells offers a model system, with their visibility greatly assisting movement analysis [22, 3]. In certain organisms, melanosomes are rapidly redistributed (see Figure 3) from aggregated (clustered near the cell’s centre) to dispersed (distributed across the cytosol): the distinct distributions alter the cell’s visibility and, when coordinated at a macroscopic level, allows the organism to change pigmentation, for example to aid camouflage [23].

**Figure 3:**
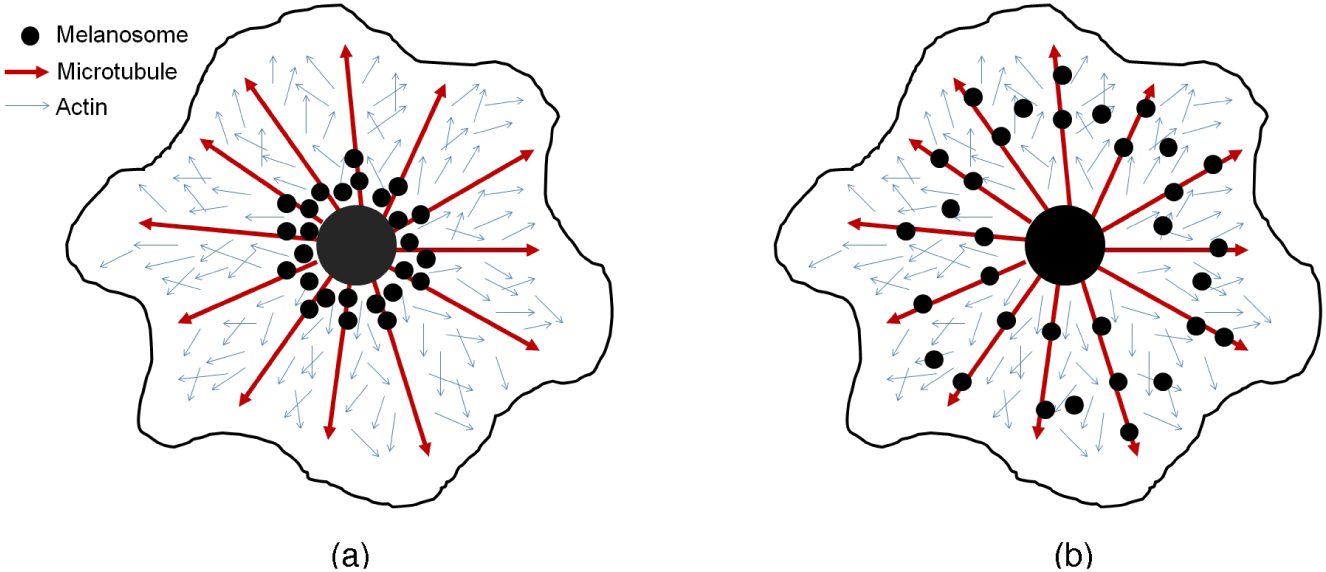
Schematic showing cytoskeletal shuttling of melanosomes in melanophores. Microtubules (red arrows) form a quasi-radial array of spokes that converge on the cell centre, with positive ends (arrow heads) pointing outwards; actin (blue arrows) forms a denser interlocking mesh, also with positive ends pointing towards the cell membrane. Melanosomes (black circles) are shuttled between an (a) aggregated state and (b) dispersed state, via motor driven movement along the microtubule and actin networks.

Organelle transport heavily relies on components of the cytoskeleton, principally the microtubule and actin networks [1]. These networks are formed by long, interlocking and polarised filament strands that grow/shrink through the addition/subtraction of proteins at the ends. Microtubules are typically fashioned into a radial array of spokes emanating from organising centres, such as the nucleus, with “positive ends” oriented outwards. Actin creates a denser mesh of short and linked fibres. Further to regulating cell shape, these networks provide a convenient transport system that allow organelles to be shifted efficiently across the cell. Movement along microtubules is achieved through organelle attachment to molecular motors that bind to microtubules, the plus-end walking kinesins and minus-end walking dyneins: the opposing directions of these motors is frequently described in the context of a “tug-of-war” [10], the balance dictating whether movement is principally outwards or inwards. Similarly, melanosomes attach to various myosin motors that walk along the actin network.

We consider a highly simplified model for melanosome shuttling along the micro-tubules, assuming that the underlying transport process is described by equation (2.5): the actin network is ignored here, but extensions to account for this and other factors will be discussed later. We assume a circular two-dimensional region, characteristic of the flattened geometry of cells in typical *in vitro* studies ([7]). The structure of the microtubule network is described by a (unimodal) von Mises distribution and is taken here to be in quasi-equilibrium over the (assumed brief) timecourse of interest: *ν*(**x**) describes the dominant direction of microtubules at position **x**, oriented according to their positive ends and with corresponding concentration parameter *k*(**x**, *t*). Taking melanosome movement to be restricted to up or down the microtubule, their turning distribution will subsequently be dictated by the bimodal von Mises distribution:

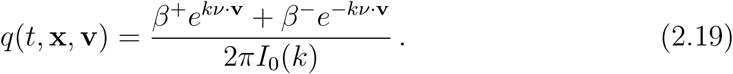

In the above, *β*^+^ and *β*^−^ (*β*^+^ + *β*^−^ = 1) respectively describe the probability of moving towards positive or negative ends; in reality, these would depend on the balance of the tethering dynein/kinesin motors, and in turn on the cell’s signalling state. Under (2.19) the macroscopic model for the melanosome distribution (*u*(*t*, *x*)) will be given by equation (2.6) with

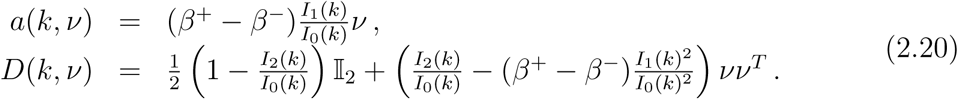

Typical microtubule networks in pigment cells are formed by radial spokes about the cell’s centre, and we therefore take *ν*(**x**) = **x**/*r* as the dominant direction and take *k*(**x**) = *k*(*r*). At the origin it is logical to assume no dominant direction exists, and therefore *k*(0) = 0. Setting *α*(*r*) = *I*_2_(*k*(*r*))/*I*_0_(*k*(*r*)) we obtain equation (1.1) with (1.2) exactly when *β*^+^ = *β*^−^: thus, the star problem emerges in the finely-poised tug-of-war where positive and negative end movement is balanced.

#### 2.2.2. Cell movement in extracellular matrix

Our next example focusses on contact-guided cell migration [5], where cells align and migrate according to the anisotropy of their environment. Contact guidance has been shown in various cells, from embryonic cells during development ([36]) to immune cells in the adult ([34]). Significantly, this behaviour is believed to facilitate the heterogeneous invasion of certain malignant cancer cells (e.g. [9, 2]). Here we focus on the impact of collagen fibres, a dominant extracellular matrix (ECM) component of stromal tissues that displays significant alignment and organisation according to tissue type [35], see Figure 4. A study in [30] indicated diverse forms of fibre arrangement surrounding implanted mammary tumours according to malignancy: in noninvasive tumours fibre orientation was predominantly parallel to the tumour boundary, effectively encapsulating the tumour; in invasive tumours, fibres became oriented orthogonal to form radial lines. In three dimensional tumour invasion assays engineered to generate these distinct arrangements, invasion depth was maximised for fibres aligned orthogonally to the tumour-matrix interface [31], consistent with a process of contact-guided orientation along fibres.

**Figure 4:**
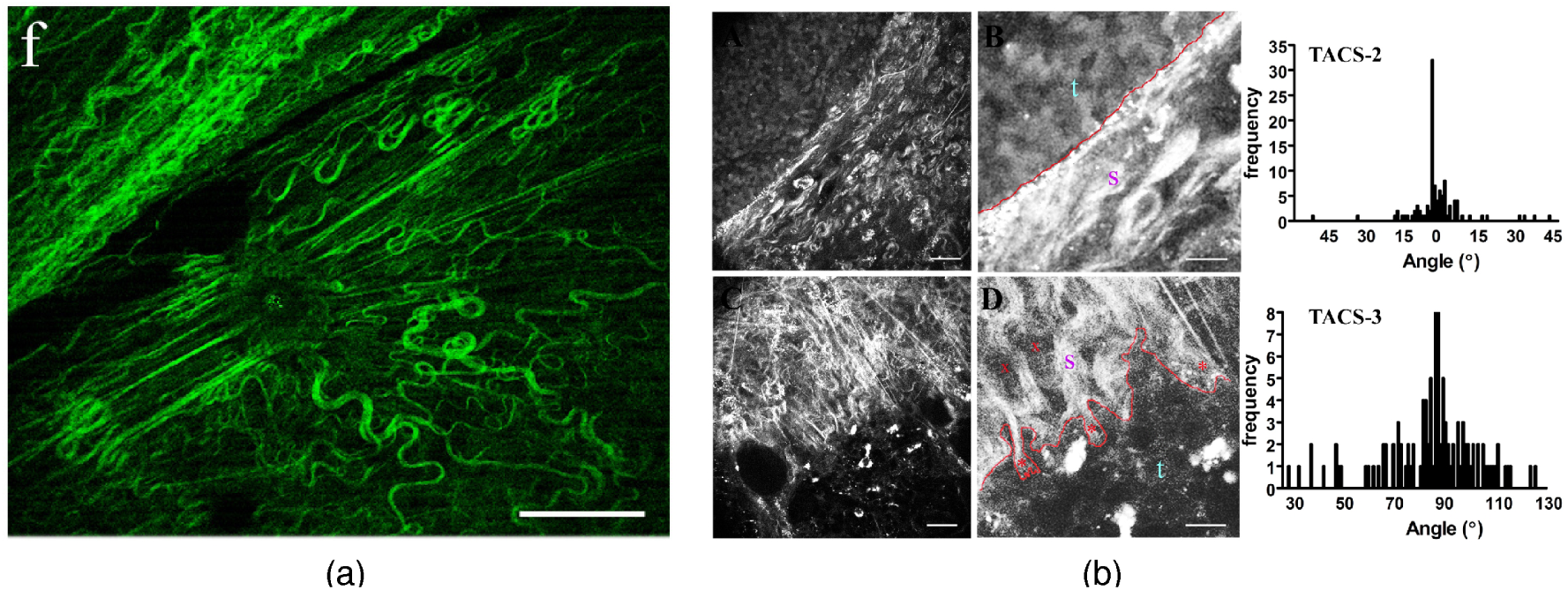
Orientation of collagen fibre networks in healthy and pathological tissues. (a) Multi-photon excitation/second harmonic generation imaging of stromal tissue surrounding the nipple, showing collagen fibers radiating from the central duct. (b) Images illustrating tumour (top-left) – matrix interfaces. (A) low magnification; (B) higher magnification of the same tumour; in this scenario, collagen is wrapped parallel to the tumour boundary (top right histogram). (C) low magnification; (D) higher magnification of the same tumour; in this scenario, collagen is aligned orthogonal to the tumour boundary (top right histogram). Tumour-stromal boundary indicated by red line, with s = stroma and t = tumour). Scale bar for A, C equals 25 *μm* and 10 *μm* for B and D. All images in this plot reproduced from Provenzano *et al.*, Collagen reorganization at the tumor-stromal interface facilitates local invasion, *BMC Medicine*, 2006, 4:38. [30].

A simplistic transport-equation based approach for contact-guided cell migration along collagen fibres was considered in [11, 26], based on equation (2.5). The anisotropic arrangement of collagen fibre networks is often quantified via bimodal von Mises distributions (e.g. see [8] and references therein), according to its (spatially-varying) dominating axial direction and alignment concentration. To model contact guidance it is therefore logical to assume cell orientation follows similarly. Specifically, we take *q* to be (2.12), where *ν* is the dominating axial direction of fibres and *k*(**x**) = *ακ*(**x**). Here, *κ*(**x**) would represent the alignment concentration parameter in the collagen fibres, while *α* is a further “tuning” parameter to reflect the fidelity of cell alignment to the fibres: larger *α* would lock cell orientation firmly along the fibre orientation, while *α* close to zero corresponds to cells that “ignore” its directional structure. Under these assumptions the equation governing the macroscopic cell density *u*(*t*, **x**) is given by equation (1.1) and (2.13).

The radial star problem subsequently corresponds to an idealised case in which fibres are radially symmetric about some origin. Particularly relevant would be the collagen fibre networks with a predominant orientation either parallel or perpendicular to the tumour/matrix boundary: under the latter we would obtain (1.3) with *α*(*r*) ∈ (−1, 0], while in the former we obtain (1.3) with *α*(*r*) ∈ [0,1).

### 2.3. Animal movement near water holes

The final example considers animal movement about watering holes. Water holes are, naturally, a prime target in arid environments and the term piosphere was coined in [18] to describe the characteristic landscape that surrounds them: principally, a central “sacrifice zone” exists where vegetation as been trampled to negligible levels and an outer zone where vegetation gradually recovers to normal (e.g. see [32, 17]). The outer zone is usually characterised by a network of well trodden radial paths, converging and merging as they center on the central zone (see Figure 5). Particular interest in piospheres stems from their impact on local flora and fauna dynamics, and subsequently determining artificial watering hole placement in agricultural (water points for livestock) and semi-wild (for example, natural parks) environments: see [32, 17]. A number of studies indicate higher animal densities in the vicinity of the watering holes: for example, aerial census surveys in Kruger national park generally reveal a trend in which herbivorous population densities decrease with radial distance from the source (representative data reproduced in Figure 5 (b)); similarly, sheep dung densities (a proxy for average population densities), have been shown to decrease with distance from artificial water sources [18] in Australian ranches.

**Figure 5:**
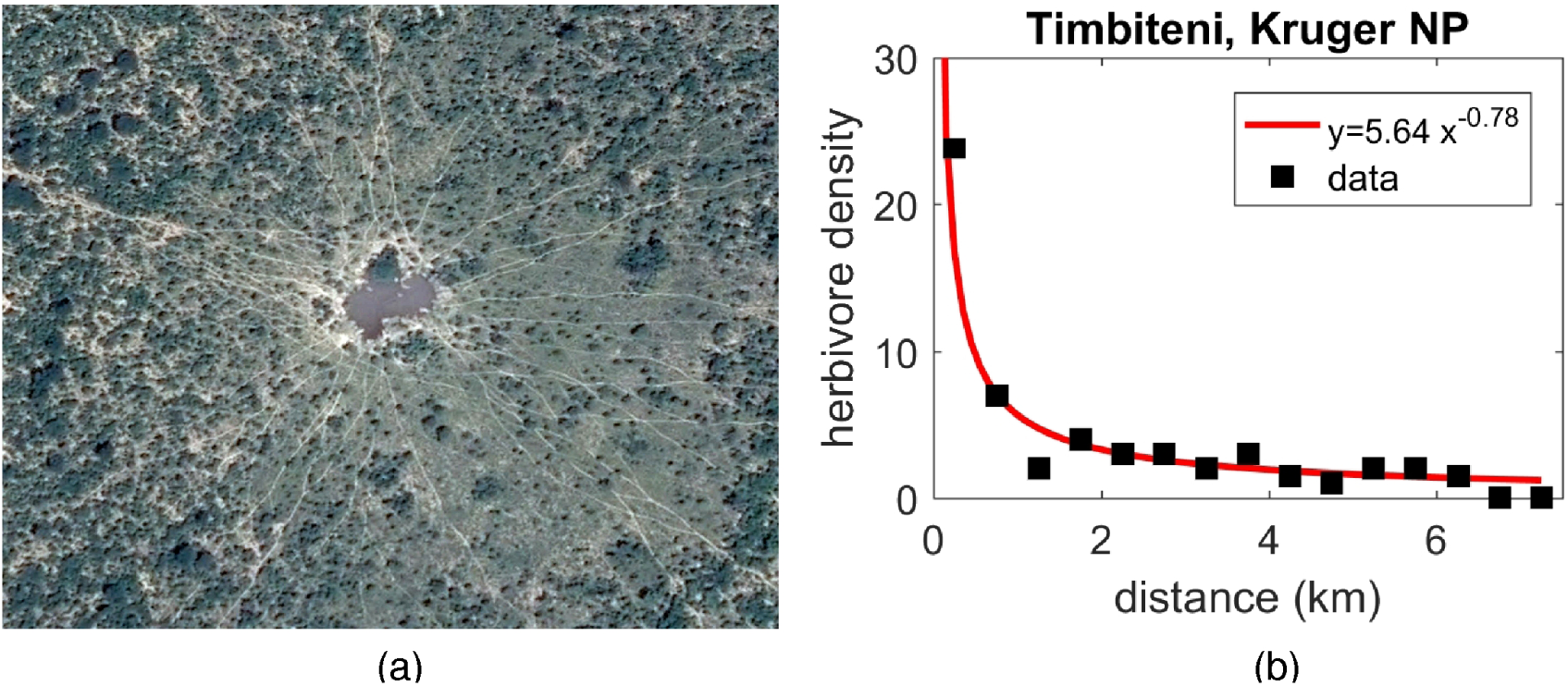
(a) A network of radial animal paths leads to a watering hole. Chobe National Park, Botswana. Image location 18°19′55″S, 24°32′04″E. Taken from Google Earth, accessed October 2016. (b) Represented data showing animal density (estimated from aerial counts) with distance from a watering hole. Data (black squares) reproduced from [33], for “Timbiteni” watering hole in Kruger National Park (South Africa). Note that herbivore density refers to “large stock units” per km^2^, a measure that takes into account each species size, resource utilisation *etc*. A power law (red line) is fitted to this dataset, see legend for coefficients.

The arrangement of animal trails leading to a waterhole offers a relevant example of (quasi) radially-symmetric orientating structure: a study of the sheep tracks emanating from a water source in an Australian ranch indicated remarkable radial alignment, typically varying by no more than a few degrees from the radial line [18]. Movement along a well-trodden trail provides the animal a path of minimal resistance and, further, limits destruction of surrounding vegetation. Taking again the transport equation (2.5) as the underlying model for movement we can once again construct the star problem through a similar set of assumptions to those for cell migration above: (i) we represent the network of trails in terms of a bidirectional von Mises distribution; (ii) we assume movement of animals along trails is bidirectional, so that the turning behaviour follows this distribution; (iii) near water holes, we assume trails to be arranged in radially-symmetric fashion that maintain an almost radial direction away from the hole (see [18]). Consequently, equation (1.1) emerges as the macroscopic model and the star problem will follow under the assumptions of radial symmetry.

### 2.4. Representative forms for α(r)

Before proceeding to analyse the star problem itself we briefly consider choices for the function *α*(*r*). The above modelling shows how *α*(*r*) can be related to the concentration parameter of the von Mises distribution: specifically, we find *α*(*r*) = ±*I*_2_(*k*(*r*))/*I*_0_(*k*(*r*)), where the sign derives from whether the dominant direction is along radial lines (+) or perpendicular to them (−). Based on the properties of *I*_2_(*k*(*r*))/*I*_0_(*k*(*r*)) and the assumptions on *k* at the origin centre, we have *α*(0) = 0 and *α*(*r*) ∈ [0,1] or [−1, 0]. We will consider the following forms:

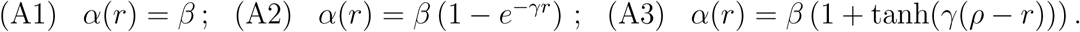

In the above, parameter *β* ∈ [0,1] for alignment along radial lines and *β* ∈ [−1, 0] for alignment perpendicular to them, while *γ*, *ρ* ≥ 0. The choice (A1) is motivated by its analytically convenient form, defining a concentration parameter that simply remains constant with distance from the origin: for *β* ≠ 0 there is a discontinuity in *α* at the origin. Form (A2) offers a smooth approximation to (A1), converging to (A1) as *γ* → ∞. The form (A3) incorporates a smooth step centred at *ρ*. This is motivated by a more plausible description of naturally occurring radial structures: rather than converging on the origin itself, microtubules radiate from the perinuclear region, collagen fibres radiate from the tumour/matrix boundary and trails emanate from the sacrifice zone surrounding a water hole. Further forms could extend these concepts further: for example, any radial alignment is likely to diminish with greater distance from the origin due to their reduced density.

## 3. Analysis of the singularity at zero

In this section we examine the structure of solutions to the star problem (1.3) under the special case of constant *α*. In particular, we systematically explore cases corresponding to *α* = 0, *α* = 1, 0 < *α* < 1, −1 < *α* < 0 and *α* = −1. In the biological examples considered earlier, a recurring element found under scenarios of positive *α* lies in its association with aggregative behaviour. For example, melanosomes can be found to be aggregated near the cell centre while surveys of animal distribution around watering holes reveal a decreasing density of animals with distance from the water hole.

### 3.1. *The isotropic case of α* = 0

In the isotropic case *α* = 0, the above model (2.18) is simply the heat equation for radial symmetric solutions in polar coordinates and subsequently the behaviour of solutions are well known. For example, if the model is equipped with homogeneous Neumann boundary conditions on a radius *R*, solutions converge to a constant solution.

### 3.2. *The fully aligned case of α* = 1

In the fully aligned case, *α* = 1, the above model (2.17) resembles a three-dimensional radial Laplace operator, but in two dimensions. We first study the steady states 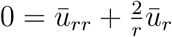. Using the ansatz *u̅* ˜ *r^β^*, we find characteristic equation

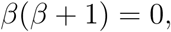

which gives two exponents: *β* = 0 and *β* = −1, respectively corresponding to a constant steady state and a non-constant steady state with a *r*^−1^ singularity at *r* = 0.

To study the time-dependent problem (2.17) we employ the scaling from the steady state analysis, i.e. we define *U*(*t*, *r*) = *ru*(*t*, *r*) and obtain for *r* > 0 that

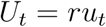

and

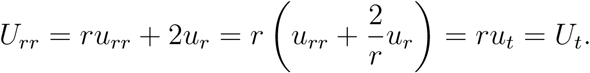

Hence, for *r* > 0 (2.17) is equivalent to:

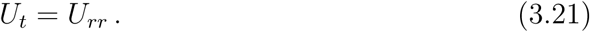

We consider this model with no-flux boundary conditions for *U* at *r* = 1:

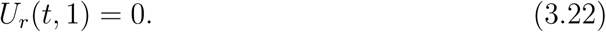

The second boundary condition stems from conservation of mass:

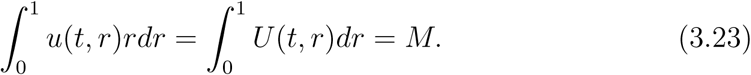

In this case we can write

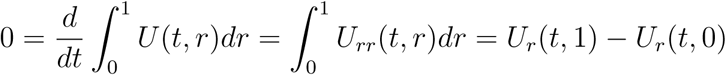

and, since *U_r_*(*t*, 1) = 0, we also have *U_r_*(*t*, 0) = 0.

The full initial-value boundary-value problem for the case *α* = 1 on a bounded radial domain is then given by

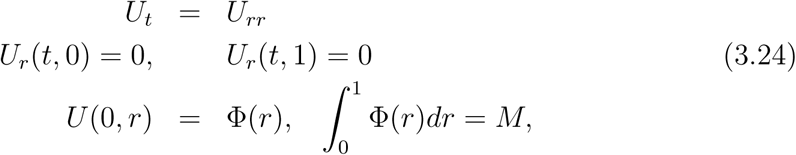

which is simply the classical heat equation with homogeneous Neumann boundary conditions, which can be solved via a classic method of separation and cosine series expansions (see, for example, introductory PDE textbooks such as [12]). For *u*(*t*,*r*) we find:

#### Theorem 3.1.

*For each* Φ ∈ *C*^0^((0,1]) *the initial-boundary value problem*

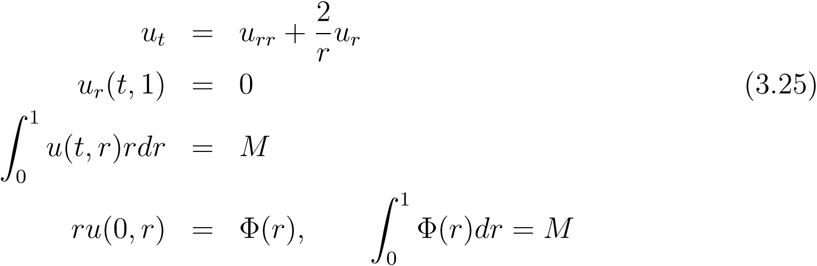

*has a unique solution u*(*t*,*r*) *with U*(*t*,*r*) = *ru*(*t*, *r*) ∈ *C*^∞^([0, ∞), *L*^2^((0,1])∩*C*^0^((0,1])). *The Fourier expansion of this solution has a leading order term M/r and the solution converges pointwise for r >* 0 *as*

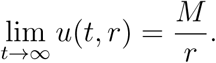

*The total mass of this solution equals M for all times t* ≥ 0.

### 3.3. *The general case of* 0 < *α* ≤ 1

For the case of constant 0 < *α* ≤ 1 we again start by studying steady states. The steady state equation for (2.16) reads

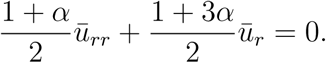

Using the ansatz *u̅*(*r*) ˜ *r^β^* we find characteristic equation

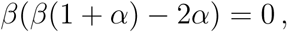

yielding exponents *β* = 0 (corresponding to constant solutions) and

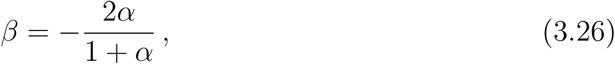

corresponding to non-constant solutions with a singularity of order 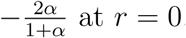 at *r* = 0.

For the time-dependent problem (2.16) we employ the scaling from the steady state analysis and define

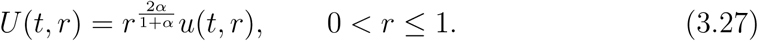

Using this transformation in equation (2.16) we obtain the following equation for *U*:

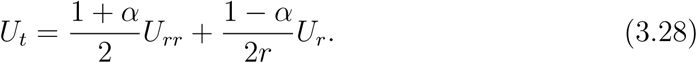

This equation is a singular Sturm-Liouville equation and we use separation of variables *U*(*r*, *t*) = *R*(*r*)*T*(*t*). The separation leads to the radial problem

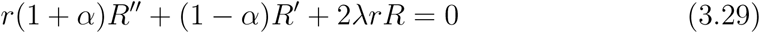

and the time problem

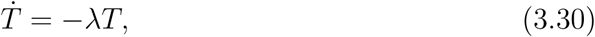

where constant λ is a Sturm-Liouville eigenvalue.

We do not solve this problem completely, rather we will only show that solutions to this Sturm Liouville problem stay bounded. First, since mass is conserved we can exclude solutions with negative λ due to their exponential growth in time (see (3.30)): hence we consider λ ≥ 0. Second, the case λ = 0 corresponds to the steady state case just studied: *U* is bounded and 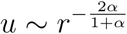.

It remains to study the case λ > 0. We define 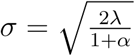 and transform variables to *s* = *σr* Then, *R*(*s*) satisfies

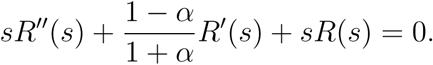

We scale *R* by an appropriate factor of *s* and write

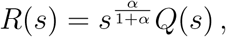

where *Q*(*s*) satisfies

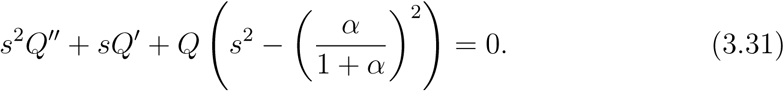

This equation is a Bessel equation of order 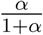. The general solution for *R* can hence be written as

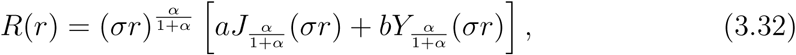

where 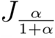 and 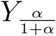 are Bessel functions of order 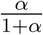 of first and second kind, respectively. The Bessel functions have the following asymptotics as *r* → 0:

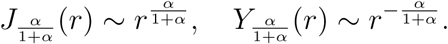

Hence all terms in (3.32) are bounded as *r* → 0. This implies that the singular behaviour near *r* = 0 corresponds to the original scaling used in (3.27), i.e.

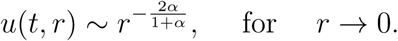

#### Theorem 3.2.

*Consider the following initial-boundary value problem for* 0 < *α* ≤ 1:

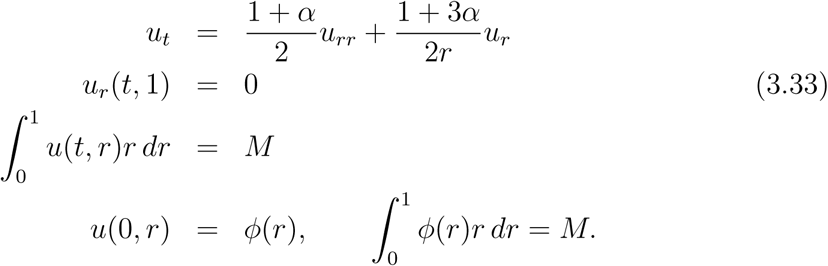

*Assume that* 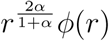 *has an absolute convergent expansion in Bessel functions of the form*

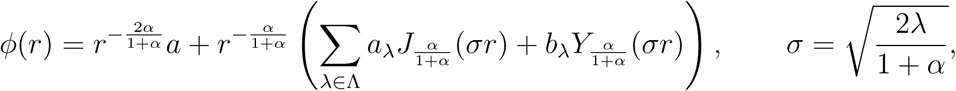

*where* Λ *is a countable set of eigenvalues. Then there exists a unique solution of the form*

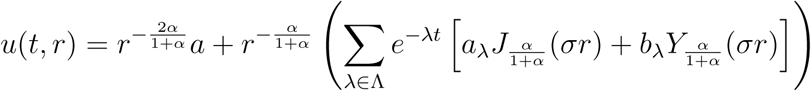

*The asymptotic behaviour near r* = 0 *depends on the coefficients a*,*a*_λ_,*b*_λ_. *If a* = 0 *and b*_λ_ = 0 *for all* λ ∈ Λ, *then the solution is bounded at r* = 0. *In all other cases*

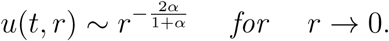

It is expected (but not proven here), that the first term 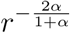 carries all the mass. In that case we can compute

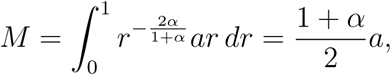

hence

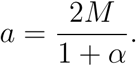

This formula makes a nice connection between the standard heat equation for *α* = 0, where *a*/2 = *M* corresponds to the first Fourier coefficient, versus the case of *α* = 1, with *a* = *M* in the fully aligned case.

### 3.4. *The case of* −1 < *α* < 0

We have noted that negative values of *α* imply rotational directions, i.e. perpendicular to the radial directions. In fact, for *α* < 0 we can perform the same calculations as above and the essential steps are summarised under introduction of positive parameter *γ* = −*α*.

The steady state problem

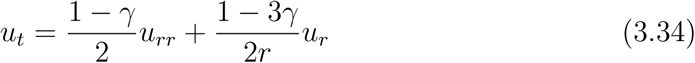

has characteristic exponents 0 and 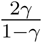. Non-constant steady state solutions therefore have the behaviour

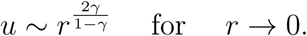

Hence, for 0 < γ < 1 the solution is continuous and equal to 0 at *r* = 0. We can use this scaling for the time-dependent problem (3.34) and consider

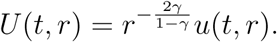

Then *U*(*t*,*r*) satisfies

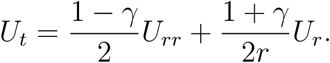

Using separation, the abbreviation 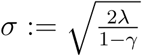 and scalings

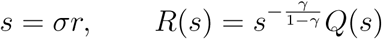

we arrive at the Bessel equation

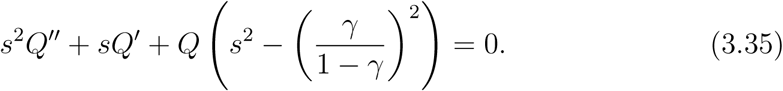

Hence the general solution for *R*(*r*) becomes

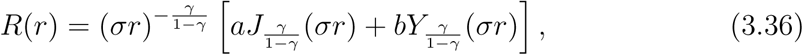

and these terms are bounded as *r* → 0.

#### Lemma 3.3.

*For the case* 0 < *α* < 1 *there is no aggregation at the origin and solutions stay bounded as r* → 0.

### 3.5. *The case of α* = −1

For the special case of perfect alignment along circles around the origin the star problem (1.3) becomes hyperbolic: 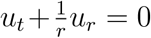. Hence, solutions are constant along characteristics 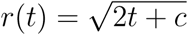, are bounded for all times and there is no aggregation at the origin.

## 4 Simulations

We numerically solve equation (1.1) along with diffusion coefficient (1.2) on a circular domain of radius *R*, subject to zero-flux boundary conditions on its circumference. Simulations are performed using COMSOL Multiphysics, which solves partial differential equation (and other) problems via finite element methods. Unless specified otherwise, initially we impose a uniformly distributed population (*u*(**x**, 0) = 1), stipulating no prior spatial structure in the population.

### 4.1. Steady state distributions and time evolution

Figure 6 plots the steady state distributions for particle populations as key parameters are changed, using the various functional forms for *α*(*r*) from (A1-A3). In particular we observe the capacity of bidirectional radial structure to organise/aggregate a population of dispersed particles, with accumulation centred on the origin. The simplest form (A1) generates a highly concentrated peak, in line with the analytical prediction of a singularity: here radial alignment of the population exists arbitrarily close to the origin, resulting in a sharply concentrated peak. Forms (A2-A3) offer regularisations, with *α*(*r*) being smooth functions that satisfy *α*(0) = 0. This regularisation translates to smooth particle profiles, although the aggregating capacity of the radial structure persists: in the case (A3) we observe a smooth profile where the particle density accumulates in the central region.

**Figure 6:**
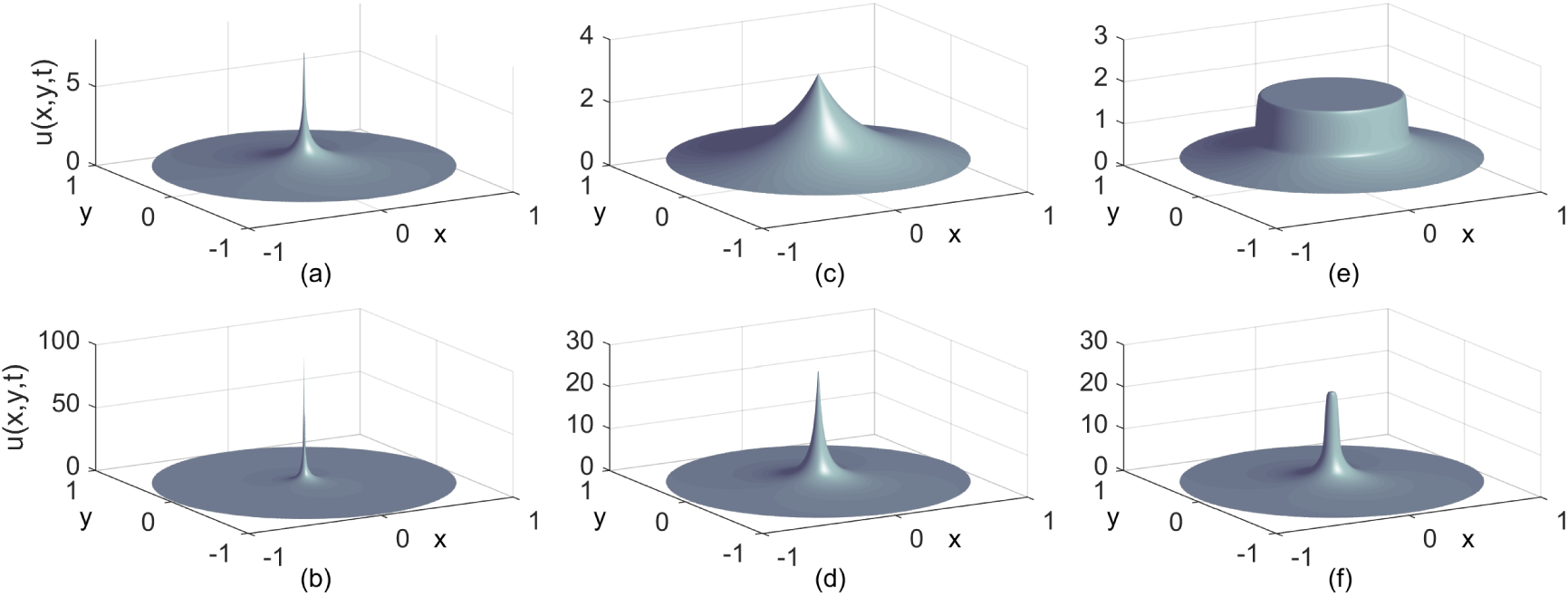
Long term (steady state) distribution of *u*(*x*,*t*) under various functional forms for *α*(*r*). (a,b) (A1) using (a) *α* = 0.25 and (b) *α* = 0.75. (c,d) (A2) using (c) γ = 1 and (d) γ = 10. (e,f) (A3) using γ = 100 and (c) ρ = 0.5 and (f) ρ = 0.05. In each case, equations (1.1-1.2) in COMSOL for a circular domain of radius *R* = 1 with zero-flux boundary conditions on the circumference, and initial conditions *u*(*x*, 0) = 1.

The solution time courses along radial lines extending from the origin are shown in Figure 7. The instantaneous singularity predicted under (A1) is revealed through the “immediate” appearance of a spike in the numerical simulations; the regularisations (A2-A3), on the other hand, demonstrate a gradual and smooth accumulation of mass at the origin.

**Figure 7:**
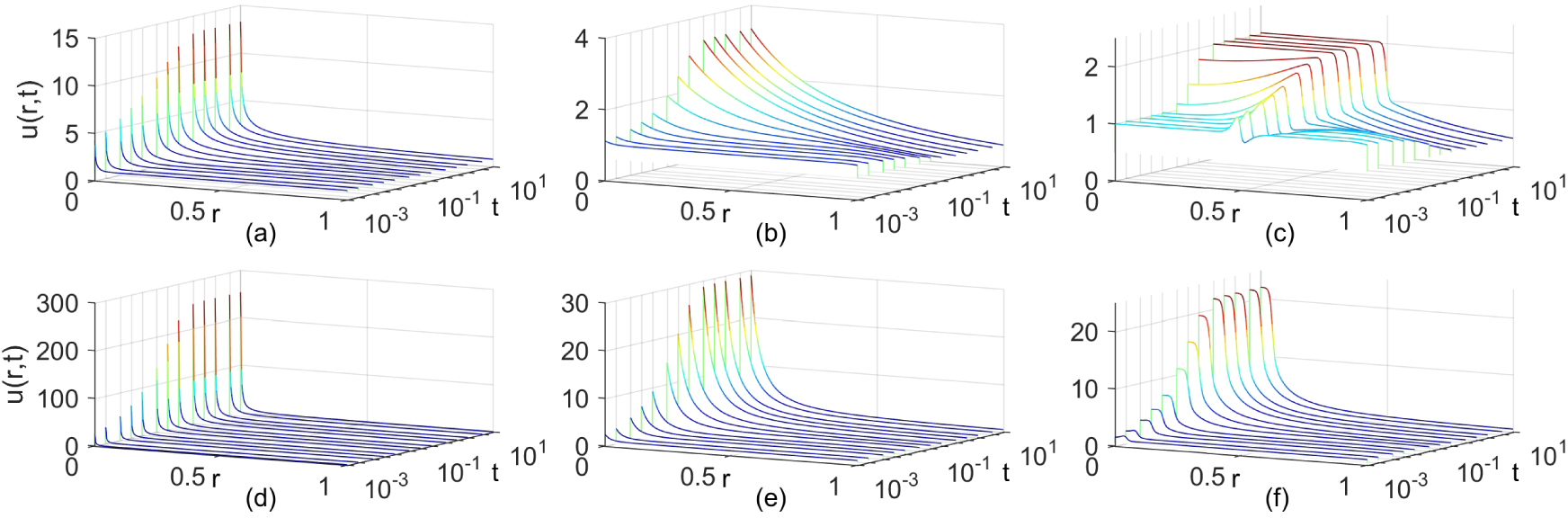
Time course showing the distribution *u*(*r*, *t*) along a radial line extending from the origin under various functional forms for *α*(*r*). (a,b) (A1) using (a) *α* = 0.25 and (b) *α* = 0.75. (c,d) (A2) using (c) *γ* = 1 and (d) γ = 10. (e,f) (A3) using *γ* = 100 and (c) *ρ* = 0.5 and (f) *ρ* = 0.05. Simulations were performed as in Figure 6, with data saved and interpolated onto a radial line extending from the origin.

We next test the validity of our leading order analytical solution for (1.3), equation (1.4), by computing this and comparing against simulated steady-state distribution for *u*(*r*, *t*). Figure 8 shows the result of this investigation: for each case considered we observe a near identical match, validating our findings.

**Figure 8:**
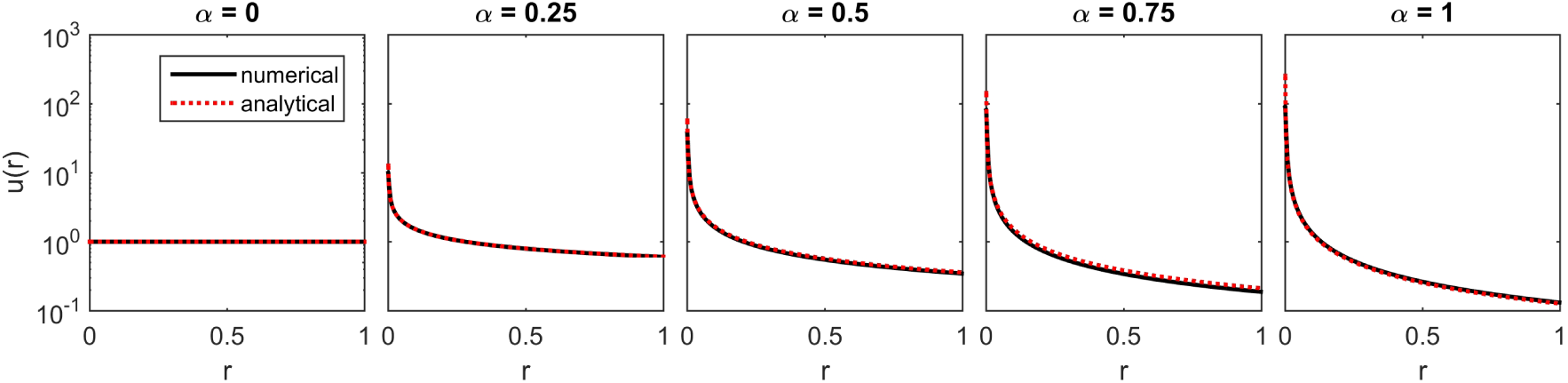
Comparison between equation (1.4) and the simulated long term steady state of equation (1.3). Numerical simulations were performed as in Figure 7.

### 4.2. Simple application: Tumour invasion patterns

The analytical and numerical results from above reveal a fundamental capacity of radial structure to dictate the movement and organisation of moving particles. We evaluate the implications in a simple application: tumour cell invasion through an *in vitro* collagen matrix assay. Our motivation comes from studies such as that of [31], where invasion assays have been devised to study how matrix anisotropy impacts on the pattern of invasion. Note, however, that our current study only explores the impact of environmental anisotropy on movement: we do not address the capacity of the invading population to itself restructure this anisotropy which, of course, could considerably influence invasion dynamics.

The configuration is illustrated in Figure 9 (a): we consider a circular assay of radius *R* and deposit an invasive cell line population near the centre. On the assay boundary we impose zero-flux conditions. We assume cells align and migrate along the collagen fibres (contact guidance) and, following our earlier arguments, utilise the simple macroscopic-level description given by (1.1) for the tumour cell density distribution *u*(*x*,*y*,*t*). The matrix surrounding the tumour cells is taken to be structured according to one of the following three forms:

1. *encapsulating* fibres arranged parallel to the tumour/matrix interface;
2. *random* fibres with no dominating orientation;
3. *radial* fibres oriented orthogonally to the tumour/matrix environment.

**Figure 9:**
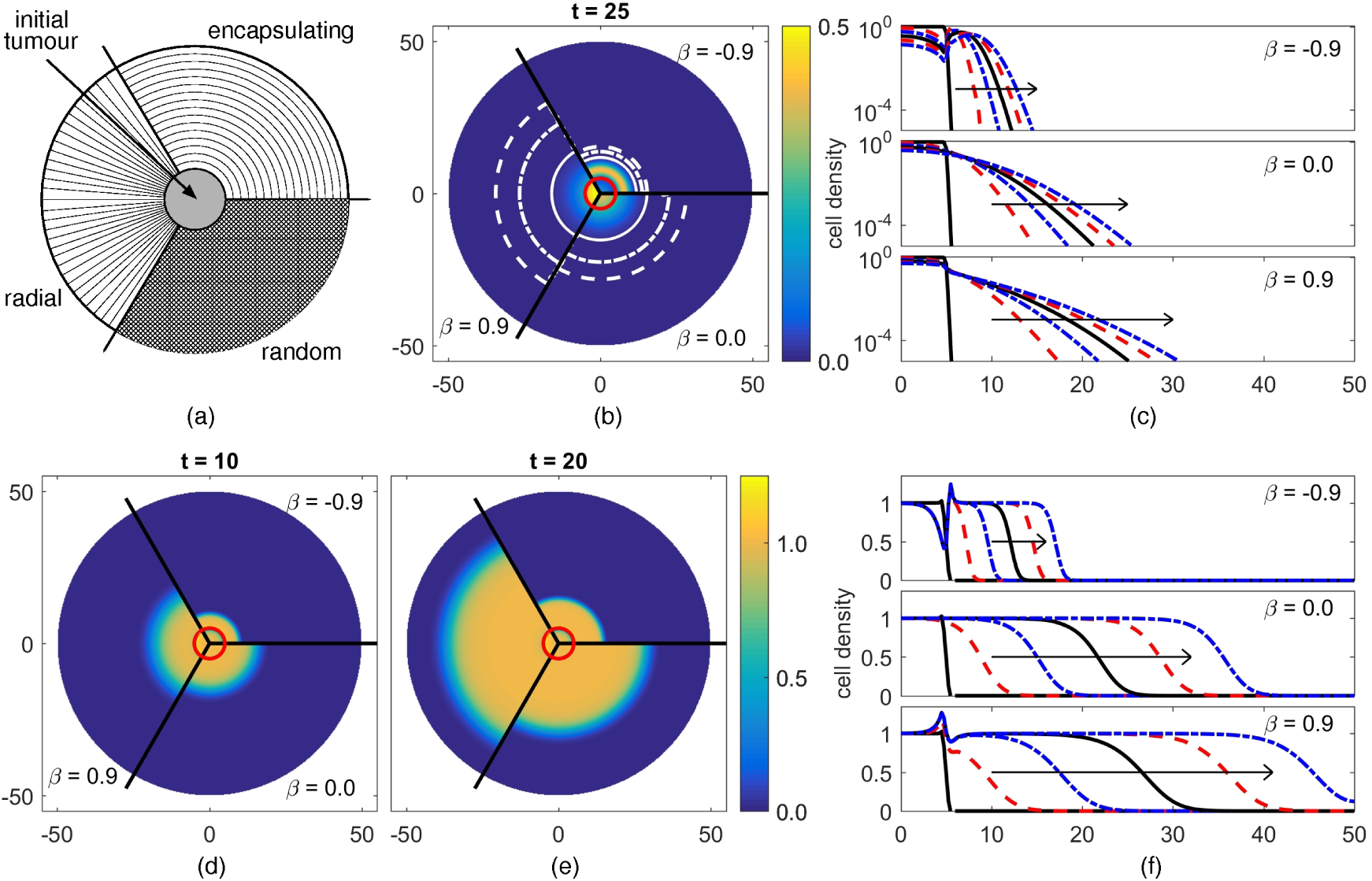
(a) Illustration of the simple tumour assay. A population of invasive tumour cells is placed in the centre of a circular assay of radius *R*. The extracellular region is composed of collagen with their predominating fibre orientation encapsulating the tumour (parallel to the tumour-matrix boundary), forming radial spokes (orthogonal) or essentially random. (b) Solution of (1.1) with (1.2) at *t* = 25 on a circular domain of radius *R* = 50. *α*(*r*) is given by (A3) with *ρ* = 5, *γ* = 100 and *β* as shown. Plot shows cell density through the included colormap, with additional contourlines marking *u* = 10^−6^ (dashed), 10^−4^ (dash-dotted) and 10^−^2. (c) Cell density plotted as a function of radius. Direction of arrows indicate increasing time, with solutions plotted at (black solid) *t* = 0, (red dashed) *t* = 5, (blue dot-dashed) *t* = 10, (black solid) *t* = 15, (red dashed) *t* = 20 and (blue dot-dashed) *t* = 25. (d-f) Solution of (1.1) and (1.2), augmented by a cell growth term *u*(1 − *u*): (d-e) Cell density profiles at times *t* = 10 and *t* = 20, (f) cell density as a function of radius; plot details as described for (b-c).

To invoke these we consider the function (A3) where we take *β* < 0 to describe encapsulating fibres, *β* = 0 to describe a random arrangement and *β* > 0 for radial fibres. We assume the central region is initially occupied by the deposited cells, with no dominating matrix orientation, so that the initial cell distribution is given by

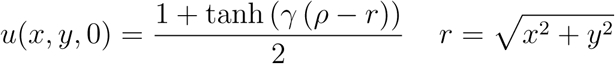

where *γ* and *ρ* take the same values as used in (A3). Note that *ρ* defines the (initial) radius of the tumour/matrix interface while *γ* is taken sufficiently large to ensure cells are initially confined within the central region.

Figure 9 (b) plots the cell density distribution at *t* = 25 for cell invasion into each of the three matrix arrangements, while in (c) profiles are plotted as a function of radius at different times; note that cell density is plotted on a logarithmic scale to highlight differences in the distribution at the leading edge of invasion. Compared to the control case of *β* = 0, an encapsulating fibre density severely hinders invasion: the population remains close to the centre with a relatively low level of infiltration taking place. Radial fibres, however, generate a double-edged effect. The aggregating nature of this arrangement, demonstrated previously, results in a large percentage of the population remaining close to the central region. This seemingly positive outcome, however, is negated by the much faster rate of infiltration at the leading edge, where a fraction of cells spread quickly outwards along the radial fibres. Consequently, radial fibre structures allow for faster infiltration by invading cells.

This effect is further illustrated via a simple extension to incorporate growth, where we augment the right hand side of equation (1.1) with a simple logistic growth term (*f*(*u*) = *ku*(1 − *u*), where *k* indicates the cell growth rate). Thus, we have Fisher’s equation under anisotropic diffusion and in a radial setting, and travelling wave solutions are observed to form. Faster infiltration along radial fibres seeds the leading edge of the wave with a greater population of tumour cells, resulting in faster invasion speeds: see Figure 9 (d-f). On the other hand, invasion is significantly limited in the case of encapsulating fibres.

## 5. Discussion

It is fascinating to observe that a purely geometric-based orientation can lead to an aggregation phenomenon in a random walk model. The aggregation effect can be strong enough that singularities arise immediately. Admittedly, the star problem represents an idealised case, and realistic biological examples will typically contain a central area that is not fully aligned. Nevertheless, we can clearly observe that aggregation can be achieved without the need for chemical chemotactic cues, adhesion processes or Turing mechanisms and arises solely from the geometric arrangement. While in the study of [15], where alignment along linear roads or parallel fibres was considered, we found convergence to *δ*-singularities in infinite time, in the star problem we can obtain immediate aggregation.

We have motivated the model in the context of three applications: melanosome transport within intracellular environments, cell migration in tissue environments and animal movement near watering holes. Although simplifying, the star problem offers insight into the role that anisotropic structure can play on population organisation.

In the context of melanosome transport, the star problem arises as a simple model for microtubule-bound movement in the harmonious case where outward (positive end/kinesin-driven) and inward (negative end/myosin-driven) movements balance: despite this parity, melanosomes naturally accumulate near the cell’s centre, replicating an aggregated state of melanosome organisation. This suggests that, in one sense, aggregation is “easier” to achieve than dispersal: for microtubule-bound melanosomes, moving into the dispersed state would require a specific bias towards outwardly directed movement. Of course our crude modelling here neglects a number of factors that would play a significant role during melanosome transport, not least the potentially crucial impact of the highly dynamic actin network on melanosome transport (e.g. see [7]). A possible avenue for further work would involve developing a more detailed model, allowing switching between movement along actin and microtubule based networks or incorporating a more sophisticated model for the molecular regulation of motor-driven transport.

Turning to animal movement, our results suggest that a simple process of path-following can generate a higher density distribution of animals close to watering holes: individuals do not require a specific movement bias towards the watering hole, rather they can simply move along a trail in bidirectional fashion and the radial convergence of paths will naturally accumulate the population at the water source. Aerial surveys in Kruger national parks reveal herbivorous population densities that decrease with radial distance from a water source (see [33], representative data reproduced in Figure 5); similarly, sheep dung densities, a proxy for average population densities, have been shown to decrease with distance from an artificial water source [18]. Significantly, the relationship between herbivore density and distance from water in these studies is usually best-fitted by power-law relationships (for example Figure 5, see also [33] and references therein), consistent with our analytical population distribution given in (1.4). Higher herbivore population densities will have a multitude of impacts on local ecology, from a manifest effect on local vegetation (creating the piosphere) to more subtle effects on predator distributions. Careful placement of artificial watering holes is therefore a conservation issue in both agricultural and natural contexts, and an obvious direction for further work would be to explore how watering hole density and distribution impacts on the environment within the context of herbivore-vegetation and/or predator-prey models.

In the context of tumour growth, our modelling suggests movement along oriented fibres can greatly facilitate invasion: a result that echoes both experimental studies (e.g. [31, 9, 2]) as well as our previous theoretical studies (e.g. [26][28]). Tumours developing from malignant cell lines can be found associated with matrix fibres oriented orthogonally to the tumour-stromal interface (e.g. [30]). In the context of the results here a radial organisation of this nature could naively be assumed to be beneficial, through aggregating the majority of the cells closer to the origin. However, those cells that do “escape” tend to infiltrate/invade much faster along radial lines for orthogonal over random or parallel fibre arrangements. Of course, we further note that invading cells will also typically modify the extracellular matrix as they invade, adding a further level of complexity to the process.

Scientific imaging in ecology and medicine is progressing quickly and we have an ever-increasing understanding of the surrounding in which species and cells exist. To understand their dynamics, the structure of the environment cannot be ignored.

## Acknowledgements

We are particularly grateful to discussions with M. Winkler during the first stages of this project. For example, he saw immediately that the 2-D star problem for *α* = 1 uses a 3-D radial Laplacian. TH’s research is supported by NSERC. KJP acknowledges the Politecnico di Torino for a visiting professorship position.

